# Deciphering the molecular mechanisms of FET fusion oncoprotein–DNA hollow co-condensates

**DOI:** 10.1101/2025.03.01.641013

**Authors:** Linyu Zuo, Qirui Guo, Kecheng Zhang, Baiyi Jiang, Zhixing Chen, Yufei Xia, Long Qian, Lei Zhang, Zhi Qi

## Abstract

Biomolecules such as nucleic acids and proteins can undergo liquid-liquid phase separation to form biomolecular condensates with diverse architectures. However, fundamental questions surrounding the biophysical principles that govern condensate architecture and their potential biotechnological applications remain unresolved. Here, we report that the FUS/EWS/TAF15 family fusion oncoprotein FUS-ERG forms hollow co-condensates with double-stranded DNA containing GGAA microsatellites. Through a combination of biochemical assays, super-resolution imaging, and mathematical modeling, we reveal that the interior surface of hollow co-condensates exhibits properties distinct from those of the external surface, a phenomenon we term nested asymmetric phase separation. Furthermore, we demonstrate that the self-organization and morphology control of FUS-ERG hollow co-condensates can be harnessed for DNA-based information manipulation, enabling targeted DNA deletion within dsDNA libraries and facilitating dynamic, hierarchical data selection. These findings provide critical insights into the biophysical mechanisms underlying multicomponent phase-separated cellular bodies and establish a foundation for leveraging condensate morphology in biotechnology.

## Introduction

Biomolecules such as nucleic acids and proteins can aggregate within living cells independently of lipid membrane encapsulation, forming mesoscale multicomponent biomolecular condensates. These condensates arise through liquid-liquid phase separation (LLPS), a critical process that facilitates their formation. LLPS-driven condensates exhibit spatiotemporal self-organization and coarsening dynamics ^1^ and are integral to numerous biological processes ^2–4^. Furthermore, dysregulation of condensate formation has been linked to the pathogenesis of various human diseases, including neurodegenerative disorders and cancer ^5^.

Multicomponent condensates exhibit a range of complex architectures, including “pearl chain”-like structures ^6–9^, nested “Russian doll” structures ^10,11^, and hollow condensate architectures ^12–14^. Among these, hollow condensates represent the simplest model that provides an entry point for studies of complex LLPS architectures. For example, Banerjee and colleagues ^13^ demonstrated the formation of hollow condensates consisting of an arginine-rich disordered nucleoprotein, protamine (PRM), in combination with RNA. They proposed that PRM and RNA assemble in a manner similar to a lipid-like diblock copolymer, where these copolymers, along with high concentrations of RNA or protein, lead to vesicle-like hollow condensates.

We report a new type of hollow condensates: one important FUS/EWS/TAF15 (FET) family fusion oncoprotein, FUS-ERG, can form hollow co-condensates with 25- base pair (bp) double-stranded DNA (dsDNA) containing a 4× GGAA microsatellite sequence (25-bp 4× GGAA dsDNA, Fig. 1b(iv)). FUS-ERG is formed by the combination of the low-complexity domain (LCD) and the RGG domain of FUS with the DNA-binding domain (DBD) of the E26 transformation-specific (ETS) family transcription factor ERG. FUS-ERG specifically binds to a microsatellite sequence characterized by GGAA repeats ^15,16^.

**Figure 1.**
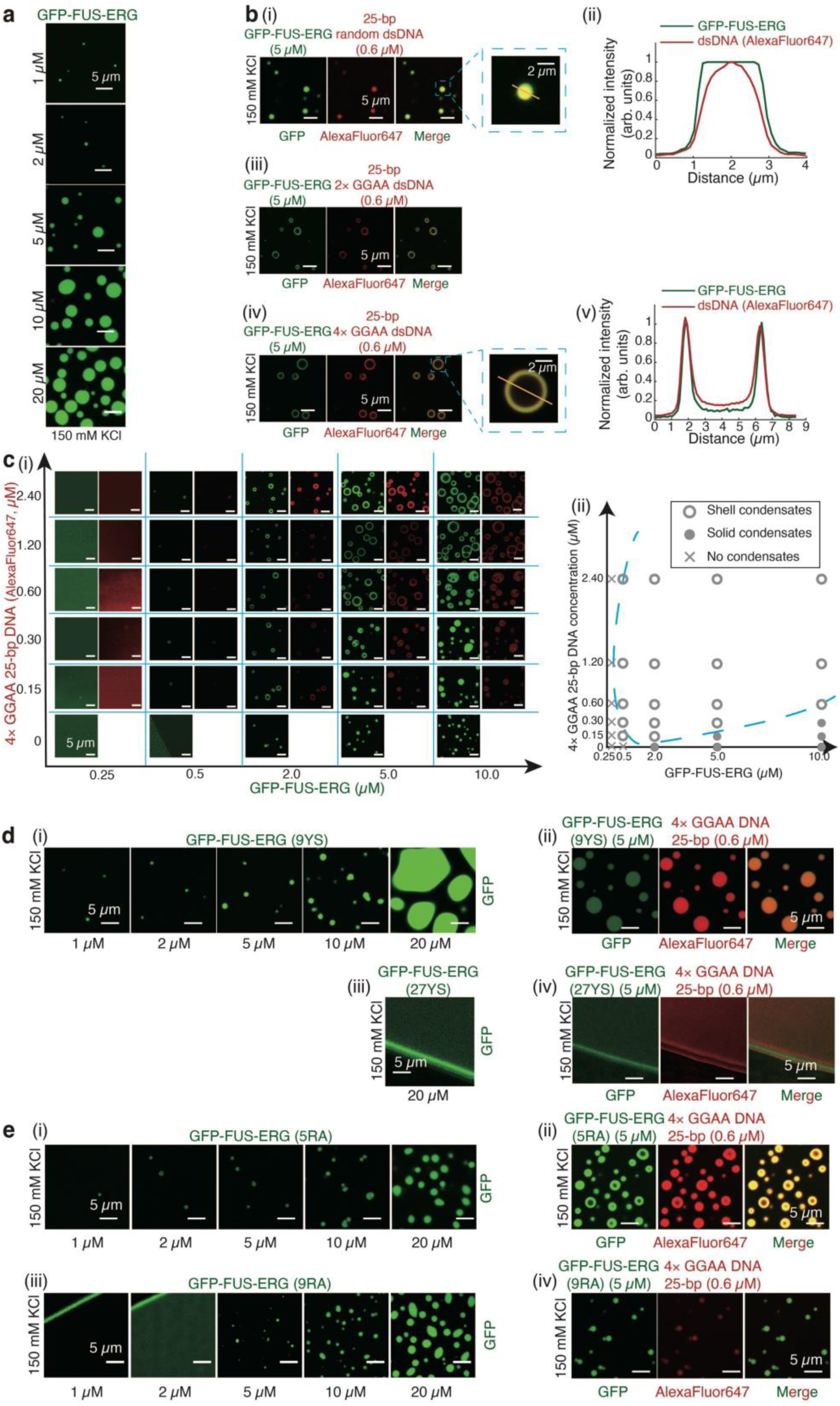
FUS-ERG can form hollow co-condensates with dsDNA containing GGAA microsatellite sequence. (**a**) GFP-FUS-ERG can undergo LLPS in the concentration of 2, 5, 10, 20 and 30 μM. (**b**) 5 μM GFP-FUS-ERG mixed with 0.6 μM 25-bp random dsDNA (i), 0.6 μM 25-bp 2× GGAA dsDNA (iii); 0.6 μM 25-bp 4× GGAA dsDNA (iv). dsDNA was labeled with AlexaFluor647. (ii) and (v) Normalized intensity profiles in (i) and (iv). (**c**) (i) GFP-FUS-ERG mixed with 25-bp 4× GGAA dsDNA; (ii) Phase diagram in (i). (**d**) (i) GFP-FUS-ERG (9YS) can undergo LLPS in the concentration of 1, 2, 5, 10, and 20 μM; (ii) 5 μM GFP-FUS-ERG (9YS) mixed with 0.6 μM 25-bp 4× GGAA dsDNA. (iii) GFP-FUS-ERG (27YS) cannot undergo LLPS in 20 μM; (iv) 5 μM GFP-FUS-ERG (27YS) mixed with 0.6 μM 25-bp 4× GGAA dsDNA. (**e**) (i) GFP-FUS-ERG (5RA) can undergo LLPS in the concentration of 1, 2, 5, 10, and 20 μM; (ii) 5 μM GFP-FUS-ERG (5RA) mixed with 0.6 μM 25-bp 4× GGAA dsDNA. (iii) GFP-FUS-ERG (9RA) can undergo LLPS in the concentration of 5, 10, and 20 μM; (iv) 5 μM GFP-FUS-ERG (9RA) mixed with 0.6 μM 25-bp 4× GGAA dsDNA. All *in vitro* droplet assays were executed under physiological conditions, specifically 40 mM Tris-HCl (pH = 7.5), 150 mM KCl, 2 mM MgCl_2_, 1 mM DTT and 0.2 mg/mL BSA, with thorough mixing and a 30-minute incubation period prior to imaging, unless otherwise indicated.

To interrogate the biophysical mechanism of FET family fusion oncoprotein hollow co-condensates and its practical implications, we employed a multidisciplinary approach. Remarkably, super-resolution imaging experiments, combined with mathematical modeling, illuminated a new molecular mechanism of nested asymmetric phase separation driving the formation of these hollow co-condensates (Fig. 4), which is distinctly different from that of vesicle-like hollow co-condensates described in previous study ^13^.

LLPS offers significant advantages in facilitating rapid biomolecular self-assembly, which has been leveraged in the design of various synthetic functional structures. For example, they were utilized to introduce compartmentalized metabolism in engineered bacteria ^17^. Because the newly discovered FET family fusion oncoprotein hollow co-condensates involve protein-dsDNA interactions, we asked whether they can provide a compelling opportunity for dynamic information manipulation in DNA-based data storage. While studies have shown that in-storage DNA encapsulation within abiotic polymers enhances data longevity and PCR uniformity, DNA encapsulation have so far been produced by pre-loading in a non-selective manner and lacked dynamic storage capacity ^18,19^. In this study, we leverage protein-DNA self-assembly based on sequence specific FET fusion oncoprotein-DNA interactions and the hollow co-condensate architecture as a dynamic and selective encapsulation medium. We demonstrated in-storage precise information separation and deletion, as well as hierarchical information sorting by altering paired DBD-dsDNA interactions. These results underscore the potential of using multi-layer LLPS for the dynamic spatial regulation of molecular information, providing a distinctive route for in-storage molecular storage and computation.

## Results

### FUS-ERG can form hollow co-condensates with dsDNA containing GGAA microsatellite sequence

We first conducted *in vitro* droplet assays to determine whether FET fusion oncoprotein FUS-ERG can undergo LLPS. We purified a GFP-tag-labeled FUS-ERG (GFP-FUS-ERG, Supplementary Fig. 1a(i)-(ii)), and performed electrophoretic mobility shift assays (EMSAs) to validate the protein activity *in vitro* (Supplementary Fig. 1a(iii)). Next, biochemical assays revealed that GFP-FUS-ERG protein can form solid condensate at a concentration as low as 1 μM *in vitro* (Fig. 1a). Mixing 0.6 μM of 25- bp dsDNA containing a random sequence (referred to as 25-bp random dsDNA) and 5 μM GFP-FUS-ERG resulted in the co-localization of both components within solid droplets (Fig. 1b(i)-(ii)). However, when dsDNA substrates of the same length were designed to contain GGAA microsatellites, such as 2× GGAA or 4× GGAA, FUS-ERG formed hollow co-condensates with these dsDNAs (Fig. 1b(iii)-(iv)). The three-dimensional hollow architecture was confirmed by confocal microscopy (Supplementary Movie 1). Fluorescence analysis revealed that 25-bp 4× GGAA dsDNA and GFP-FUS-ERG co-localized on the shell of hollow co-condensates (Fig. 1b(v)). Interestingly, although 25-bp random dsDNA and GFP-FUS-ERG could also co-localize to form solid condensates, GFP-FUS-ERG condensates enveloped the 25- bp random dsDNA (Fig. 1b(ii)), suggesting that the protein itself has stronger interactions with the solvent. Remarkably, when the length of random dsDNA increased from 25-bp to 306-bp, hollow co-condensates can also be formed, with a larger diameter compared to that of 25-bp random dsDNA (Supplementary Fig. 2).

We investigated the conditions required for hollow co-condensate formation through *in vitro* droplet assays, combining 25-bp 4× GGAA dsDNA at concentrations of 0, 0.15, 0.3, 0.6, 1.2, and 2.4 μM with GFP-FUS-ERG protein at concentrations of 0.25, 0.5, 2, 5, and 10 μM (Fig. 1c(i)). The resulting phase diagram (Fig. 1c(ii)) reveals that the DNA-to-protein molar ratio beyond a threshold number ([DNA] / [protein] ∼ 0.06) drives the hollow co-condensate formation.

Both hollow and solid condensates exhibited slow fusion kinetics (Supplementary Fig. 3a). Following this, we conducted fluorescence recovery after photobleaching (FRAP) experiments (Supplementary Fig. 3b(i)-(ii) and **Methods**) using GFP-FUS-ERG alone or mixed with 25-bp 4× GGAA dsDNA. The molecular dynamics of GFP-FUS-ERG in hollow co-condensates were slower compared to solid condensates, suggesting a more gel-like material property inside the shell region of hollow co-condensates.

While modifying the dsDNA sequence and length has been shown to regulate the formation of hollow condensates, we next asked how protein can affect the condensate morphology. We first focused on the LCD of FUS-ERG. The FUS LCD contains 27 [G/S]Y[G/S] repeats, and these tyrosines are important for FUS’s phase separation capacity ^20,21^. When 9 relevant tyrosine residues were changed to serine in GFP-FUS-ERG (9YS) (Supplementary Fig. 1c), the phase separation capacity remained unchanged (Fig. 1d(i)). However, 5 μM GFP-FUS-ERG (9YS) cannot form hollow co-condensates with 25-bp 4× GGAA dsDNA (Fig. 1d(ii)). When all 27 tyrosines were mutated to serine in GFP-FUS-ERG (27YS) (Supplementary Fig. 1d), this mutant lost its phase separation capacity even at 20 μM (Fig. 1d(iii)), and 5 μM GFP-FUS-ERG (27YS) with dsDNA also failed to form droplets (Fig. 1d(iv)).

Second, we investigated the role of the RGG motif in FUS-ERG. Mutation of five key arginine residues to alanine (GFP-FUS-ERG (5RA), Supplementary Fig. 1e) did not affect the phase separation capacity (Fig. 1e(i)). In contrast, mutating nine arginine residues to alanine (GFP-FUS-ERG (9RA), Supplementary Fig. 1f) significantly reduced the phase separation capacity (Fig. 1e(iii)). GFP-FUS-ERG (5RA) formed both solid and hollow co-condensates with 25-bp 4× GGAA dsDNA (Fig. 1e(ii)). Conversely, GFP-FUS-ERG (9RA) only formed smaller solid condensates with 25-bp 4× GGAA dsDNA (Fig. 1e(iv)). GFP-FUS-DDIT3, which is another FET fusion oncoprotein, has been reported to form similar spherical shell *in vivo* ^22^. We repeated this experiment in U2OS cells (Supplementary Fig. 4a). Interestingly, when all arginine residues in the RGG motif were mutated to alanine, FUS-DDIT3-GFP (9RA) also cannot form this architecture *in vivo* (Supplementary Fig. 4b). Taken together, these results indicate that the LCD and RGG motifs in FUS-ERG can regulate the hollow co-condensate formation.

To further elucidate the role of dsDNA and protein, we designed a two-step experiment. Initially, we used GFP-FUS-ERG to form solid condensates (Fig. 1a). Subsequently, we introduced 25-bp 4× GGAA dsDNA (Fig. 2a(i), time 0). Remarkably, the dsDNA substrates promptly enveloped the exterior surface of the solid GFP-FUS-ERG condensates (Fig. 2a(v)). This process was monitored continuously for 90 minutes (Fig. 2a and Supplementary Movie 2). By time 30-minute mark (Fig. 2a(ii)), the dsDNA substrates had completely infiltrated the condensate, coinciding with a reduction in protein intensity within the central region (Fig. 2a(vi)). By the 90-minute timepoint (Fig. 2a(iv)), hollow co-condensates had fully formed, with both dsDNA and protein concentrated on the shell of the hollow co-condensates (Fig. 2a(vii)). As a control experiment, when 25-bp random dsDNA was used, no hollow co-condensates formed even after 90 minutes (Fig. 2b), and significantly fewer dsDNA molecules were transferred into the GFP-FUS-ERG condensates (Fig. 2c). These findings demonstrate the spontaneous transfer of dsDNA containing GGAA microsatellites into the condensates accompanying hollow co-condensate formation.

**Figure 2.**
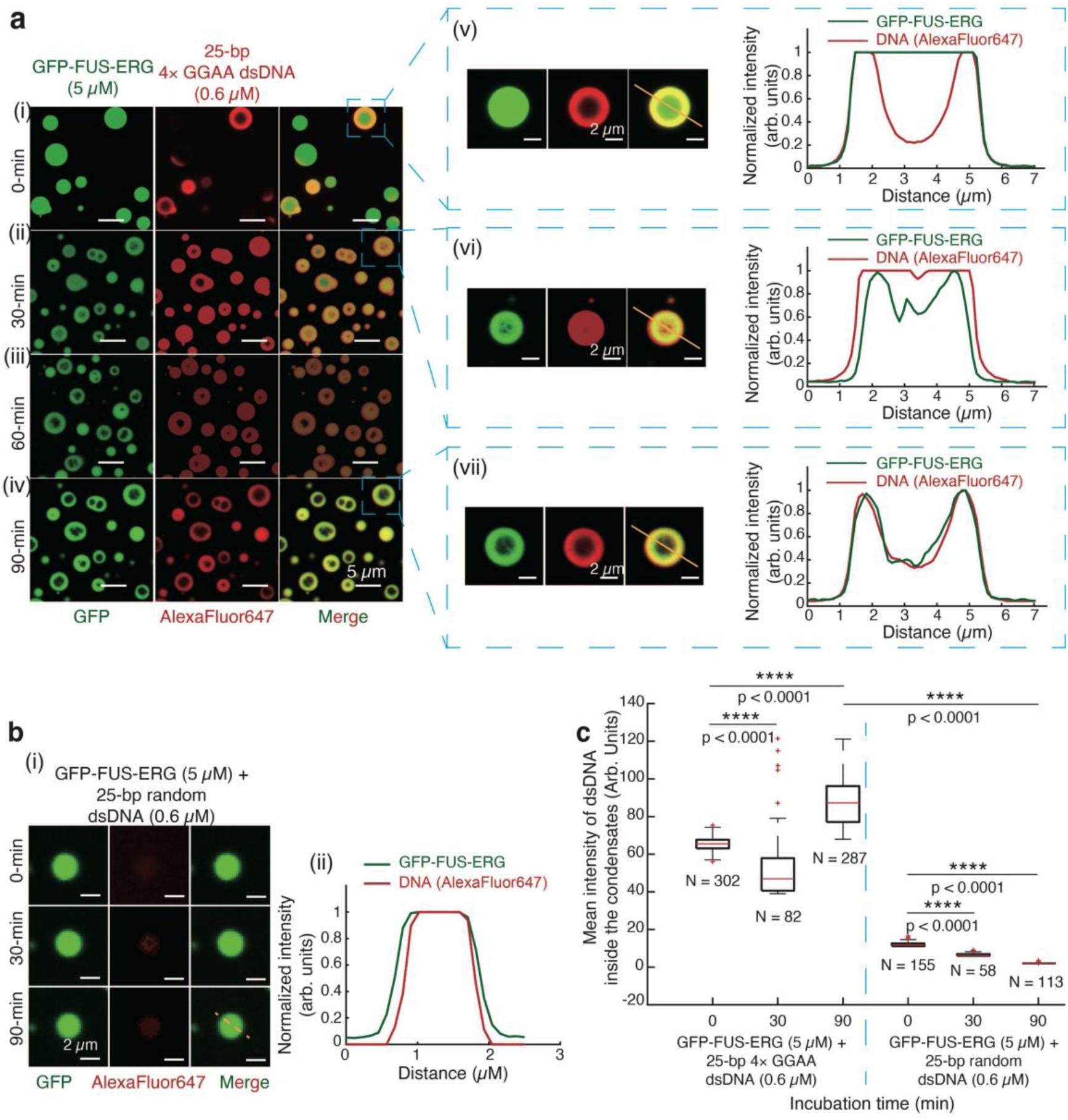
25-bp dsDNA substrates containing GGAA microsatellites can transfer into the solid FUS-ERG condensates, inducing the hollow co-condensate formation. (**a**) Time course of hollow co-condensate formation of 5 μM GFP-FUS-ERG mixed with 0.6 μM AlexaFluor647-labeled 25-bp 4× GGAA dsDNA at 0-min (i), 30-min (ii), 60-min (iii), and 90-min (iv). At the 0-minute time point, we injected dsDNA. (v), (vi), and (vii) are representative events of hollow co-condensate formation and normalized intensity profiles from (i), (ii), and (iv). (**b**) (i) Time course of hollow co-condensate formation of 5 μM GFP-FUS-ERG mixed with 0.6 μM AlexaFluor647- labeled 25-bp random dsDNA at 0-min, 30-min, and 90-min. At the 0-minute time point, we injected dsDNA. (ii) is the normalized intensity profile at the 90-minute time point. (**c**) Boxplot of the mean intensity of dsDNA inside the condensates for GFP-FUS-ERG with 25-bp random dsDNA and 25-bp 4× GGAA dsDNA. The total number N examined over one-time *in vitro* droplet experiments. For the boxplot, the red bar represents median. The bottom edge of the box represents 25^th^ percentiles, and the top is 75^th^ percentiles. Most extreme data points are covered by the whiskers except outliers. The ‘+’ symbol is used to represent the outliers. Statistical significance was analyzed using unpaired t test for two groups. P value: two-tailed; p value style: GP: 0.1234 (ns), 0.0332 (*), 0.0021 (**), 0.0002 (***), <0.0001 (****). Confidence level: 95%.

### The formation of hollow co-condensates is driven by nested asymmetric phase separation

To uncover the molecular mechanism driving the formation of hollow co-condensates, we focused on the processes that define the development of both the external and internal surfaces. GFP-FUS-ERG alone undergoes LLPS with the working buffer (Fig. 1a), suggesting that the stable external surface arises from interactions between free proteins and the buffer. Furthermore, in co-condensates formed by GFP-FUS-ERG and 25-bp random dsDNA, the GFP-FUS-ERG boundary extended beyond the dsDNA boundary (Fig. 1b(i)-(ii)). This observation strongly indicates that the external surface of the hollow co-condensates results from phase separation between free proteins and the surrounding buffer.

To elucidate the formation of the internal surface, we initiated an investigation employing RNA as a probe to discern the characteristics of the hollow co-condensates. Specifically, we combined SNAP-tag-labeled FUS-ERG (SNAP-FUS-ERG, Supplementary Fig. 1b(i)-(iii)) with 25-bp 4× GGAA dsDNA to induce the formation of hollow co-condensates (Supplementary Fig. 1b(iv)). After a 30-min incubation with Poly-U RNA, we observed the penetration of the RNA substrates through the shell, resulting in their enrichment within the lumens of the hollow co-condensates (Fig. 3a and d). Control experiments were conducted to corroborate these findings. Specifically, we observed that RNA substrates encountered difficulty in penetrating the solid condensates formed in conjunction with 25-bp random dsDNA (Fig. 3b and d) or by SNAP-tag-labeled FUS-ERG alone (Fig. 3c-d).

**Figure 3.**
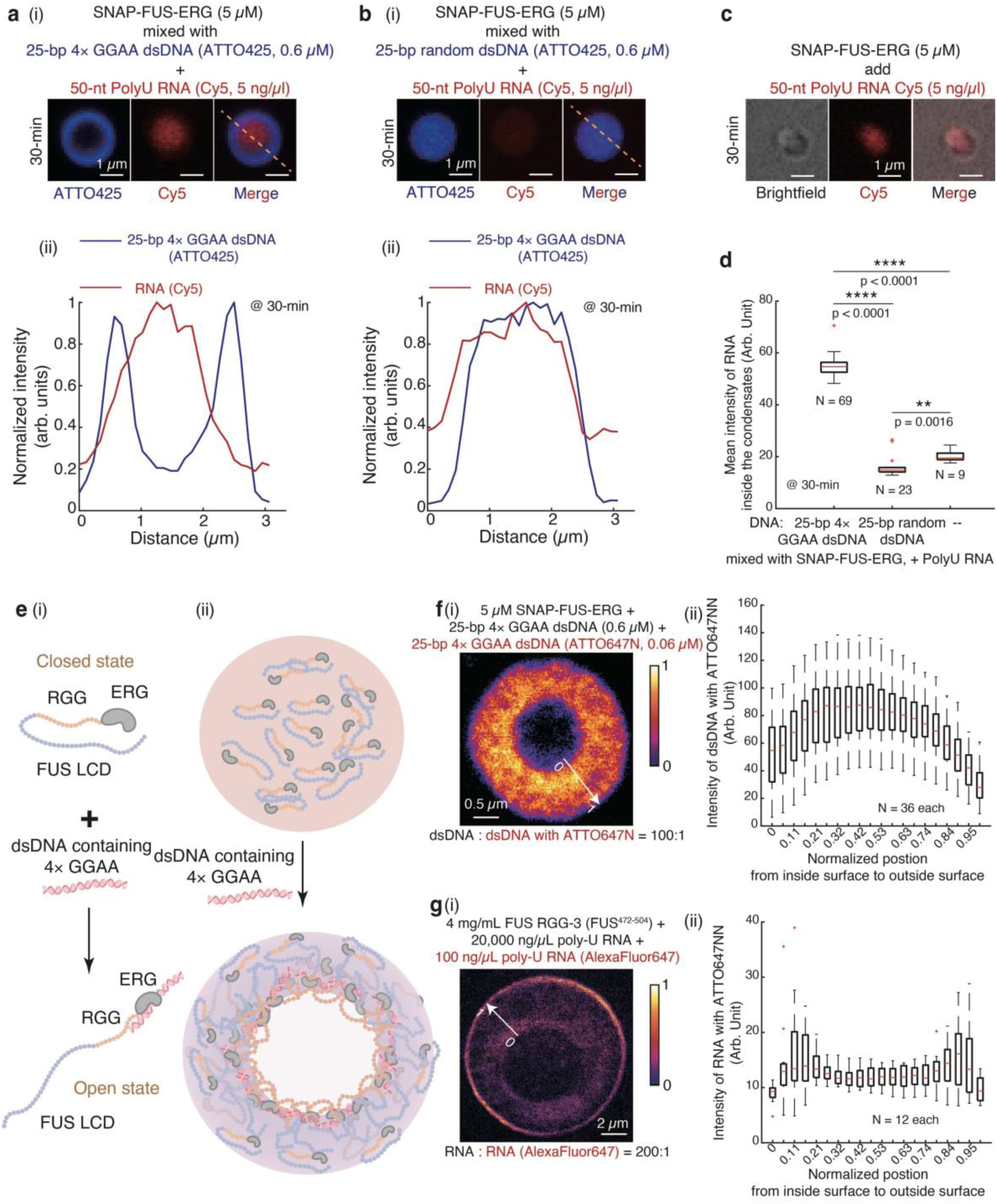
RNA probe and STED imaging reveal that the formation of hollow co-condensates is driven by nested asymmetric phase separation. (**a-c**) 5 μM GFP-FUS-ERG first mixed with 0.6 μM AlexaFluor647-labeled 25-bp 4× GGAA dsDNA (a(i)), 0.6 μM AlexaFluor647-labeled 25-bp random dsDNA (b(i)), or no dsDNA (c). 5 ng/μL 50-nt Cy5-labeled PolyU RNA was injected in the second step. a(ii) and b(ii) are normalized intensity profiles from a(i) and b(i). (**d**) Boxplot of the mean intensity of RNA inside the condensates in a(i), b(i) and c. The total number N examined over one-time *in vitro* droplet experiments. (**e**) Schematics of possible conformational change of FUS-ERG and dsDNA containing 4× GGAA (i) and hollow co-condensate formation. (**f**) (i) Representative fluorescence heatmap of STED image for 5 μM SNAP-FUS-ERG mixed with 25-bp 4× GGAA dsDNA: 0.6 μM dark dsDNA and 0.06 μM dsDNA labeled with Atto647N. The molar ratio is 100:1. (ii) Boxplot of the radial intensity distribution of dsDNA labeled with Atto647N within the hollow co-condensates. The total number N examined over one-time *in vitro* droplet experiments. (**g**) (i) Representative fluorescence heatmap of STED image for 4 mg/mL FUS-RGG3 (FUS No. 471-504) mixed with Poly-U RNA: 20,000 ng/μL dark RNA and 100 ng/μL RNA labeled with AlexaFluor 647. The ratio is 200:1. (ii) Boxplot of the radial intensity distribution of RNA labeled with AlexaFluor 647 within the hollow co-condensates. The total number N examined over one-time *in vitro* droplet experiments. For the boxplot in d, f(ii) and g(ii), the red bar represents median. The bottom edge of the box represents 25^th^ percentiles, and the top is 75^th^ percentiles. Most extreme data points are covered by the whiskers except outliers. The ‘+’ symbol is used to represent the outliers. Statistical significance was analyzed using unpaired t test for two groups. P value: two-tailed; p value style: GP: 0.1234 (ns), 0.0332 (*), 0.0021 (**), 0.0002 (***), <0.0001 (****). Confidence level: 95%. The working buffer containing 40 mM Tris-HCl (pH 7.5), 150 mM KCl, 2 mM MgCl_2_, 1 mM DTT and 0.2 mg/mL BSA.

The RGG motif (amino acids 485–539) (Supplementary Fig. 1a(i)) is the only domain within GFP-FUS-ERG capable of tightly binding RNA. These findings strongly suggest that the interaction with dsDNA containing GGAA motifs induces a conformational change in FUS-ERG (Fig. 3e(i)). In the absence of dsDNA, the RGG motif is likely sequestered within the protein, resulting in weak RNA binding and a conformation referred to as the Closed state. However, upon binding to 25-bp 4× GGAA dsDNA, the protein-dsDNA interaction appears to trigger a conformational change that exposes the RGG motif, transitioning the protein to an Open state (Fig. 3e(i)).

Integrating the RNA probe experiments (Fig. 3a-d) with earlier observations demonstrating the translocation of GGAA motif-containing dsDNA into the condensates (Fig. 2a) offers an interesting hypothesis for the formation of the internal surface. When a dsDNA substrate containing GGAA motifs binds to a free FUS-ERG molecule near the condensate boundary (Fig. 2a(i)), a protein–dsDNA complex is formed, inducing a conformational change in the protein from the Closed state to the Open state (Fig. 3e(i)). The results shown in Fig. 2 suggest that this protein–dsDNA complex can translocate into the condensate’s inner region, where the exposed hydrophilic RGG motif (Supplementary Fig. 5a) accumulates, thereby establishing the internal surface (Fig. 3e(ii)).

If this hypothesis holds true, the interior surface of the hollow co-condensates is expected to exhibit the highest concentration of dsDNA–protein complexes, with a gradual radial decrease in concentration toward the exterior. Consequently, dsDNA substrates are anticipated to display an asymmetric distribution between the interior and exterior surfaces. To investigate this distribution, we employed Stimulated Emission Depletion (STED) microscopy, which provides sub-100 nm resolution ^23^, to image the hollow condensates. For optimal brightness, dsDNA substrates were labeled with ATTO647N at a 1:100 ratio. Fluorescent signals from the labeled dsDNA substrates are presented in Fig. 3f(i). Quantitative imaging analysis (**Supplementary Methods**) revealed that the interior surface exhibited significantly higher dsDNA intensity, with a gradual decrease in intensity along the radial axis toward the exterior (Fig. 3f(ii)), confirming the asymmetric distribution of the FUS-ERG–dsDNA complex.

The hollow co-condensates formed by PRM and RNA ^13^, are expected to exhibit a symmetric distribution of RNA due to their vesicle-like characteristics. As the RGG motifs of FUS is similar to the PRM in RNA binding abilities, we repeated the STED experiment using the third RGG motif inside FUS (FUS RGG-3, 471-504) and Poly-U RNA, and obtained consistent results (Supplementary Fig. 6). The fluorescent signals of RNA substrates are depicted in Fig. 3g(i), and subsequent image analysis confirmed the symmetric distribution of RNA substrates (Fig. 3g(ii)). Collectively, these super-resolution imaging results elucidate a new type of molecular mechanisms underlying the internal surface formation of FUS-ERG and dsDNA hollow co-condensates, which we term “nested asymmetric phase separation”.

### A mathematical model not only reproduces the formation of hollow co-condensates but also predicts their selective capacity for DNA

To elucidate the molecular mechanism underlying hollow co-condensate formation, we developed a molecularly informed phase-field model ^24^. This model incorporates three key order parameters. The first order parameter, *η* = *ζ*_*hydrophilic*_ + *ζ*_*hydrophobic*_ − *ψ*_*C*_, represents the concentration of protein–dsDNA complexes (Open state in Fig. 3e(i)) within the hollow co-condensates. Here, *ζ* denotes the component concentration (volume fraction), with *ζ*_*hydrophilic*_ and *ζ*_*hydrophobic*_ corresponding to the hydrophilic and hydrophobic concentrations of the protein–dsDNA complex, respectively. ψ_*C*_ is the critical volume fraction required for phase separation. The second order parameter, *ϕ* = *ζ*_*hydrophilic*_ − *ζ*_*hydrophobic*_, indicates local hydrophilicity (positive values) or hydrophobicity (negative values) within the hollow co-condensates. Using *η* and *ϕ*, we derived the third order parameter, *χ*, which represents the dsDNA concentration within the hollow co-condensates. The free energy functional of this mesoscopic model was constructed following the classic Ohta–Kawasaki framework ^25,26^. Detailed descriptions of the model development and numerical simulations are provided in the **Supplementary Methods**.

Fig. 2a(i)-(ii) shows that dsDNA molecules containing GGAA microsatellites initially localize at the surface of FUS-ERG condensates before gradually diffusing into the condensate interior. These observations informed the initial conditions of our model (Fig. 4a(i)), where the concentrations of protein and dsDNA are denoted as *ζ*_*DNA*_ and *ζ*_*protein*_, respectively, to accurately replicate the experimental data. The diffusion-driven simulation, which considers only dsDNA diffusion, revealed a gradual inward migration of dsDNA molecules into the condensates (Fig. 4a(ii)), closely aligning with the experimental results shown in Fig. 2a(ii).

**Figure 4.**
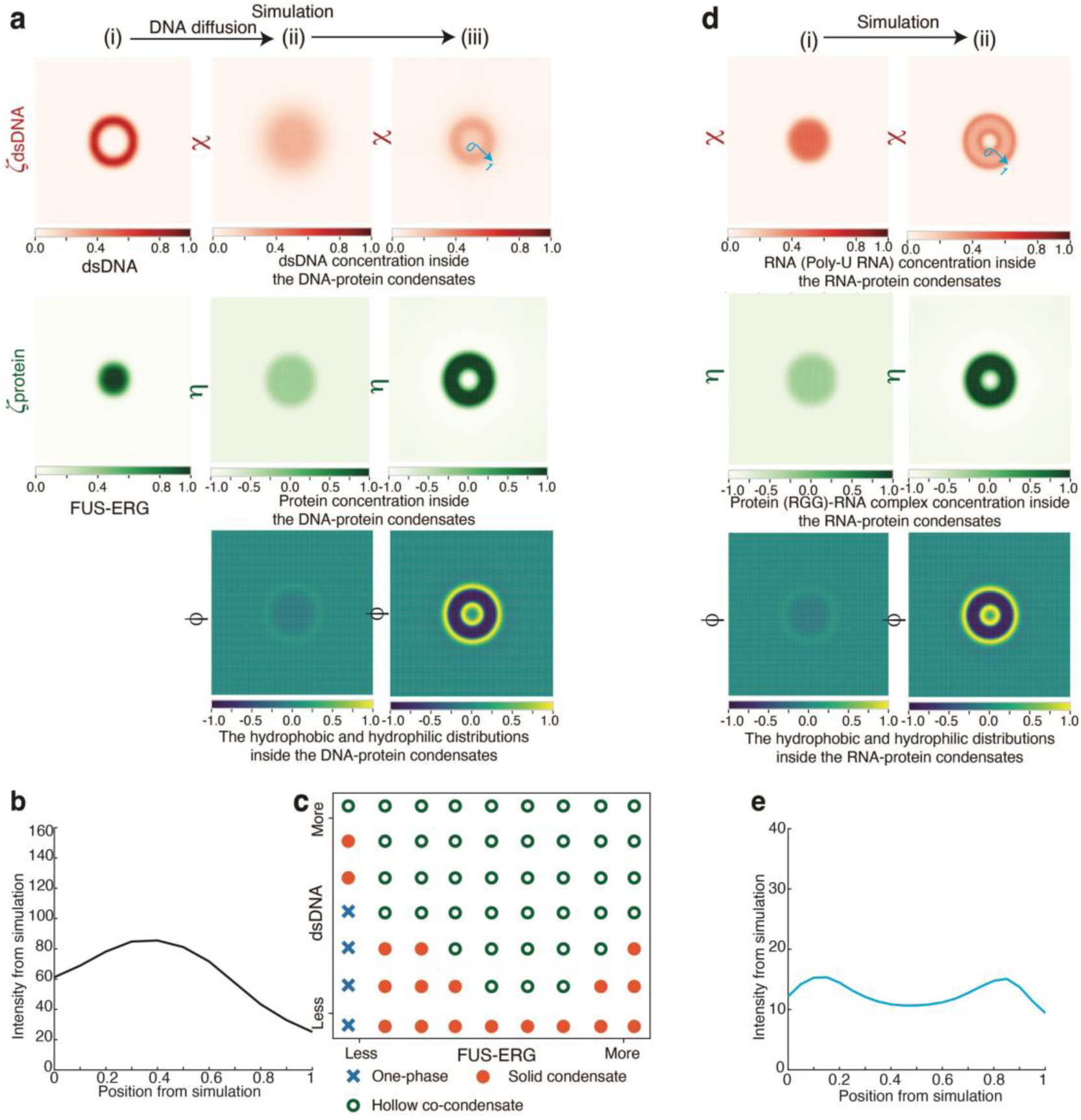
A mesoscopic, molecularly-informed phase field model was developed to reproduce the hollow co-condensate formation. (**a**) Simulations for FUS-ERG- DNA hollow co-condensate formation. (i) FUS-ERG molecules form biomolecular condensates with dsDNA containing GGAA microsatellites on their surface. dsDNA (*ζ*_*DNA*_) and protein (*ζ*_*protein*_) concentrations are represented. (ii) dsDNA molecules are transited into protein droplets, triggering the formation of hollow co-condensates. Order parameters denoting protein-DNA complex concentration (η), dsDNA concentration (χ) and hydrophobic and hydrophilic distributions within condensates (ϕ) are represented. (iii) Hollow architecture is observed in steady states of the model proposed. DNA is coupled with hydrophobic region of the protein-DNA complex. (**b**) Distributions of simulated dsDNA concentration (χ) within hollow architectures. (**c**) States diagram of simulated structures by varying initial states in a(i). Blue cross: One-phase, which is a homogeneous state; Orange dot: solid condensates; Green circle: hollow co-condensates. (**d**) Simulations for PRM-RNA hollow co-condensate formation. (i) RNA first forms tadpole-like diblock copolymer with PRM protein. Order parameter denoting protein-RNA complex concentration (η), RNA concentration (χ) and hydrophobic and hydrophilic distributions within copolymer (ϕ) are represented. (ii) Hollow structure is observed in simulation results. RNA is coupled with both hydrophobic and hydrophilic region of protein-RNA complex. (**e**) Distributions of simulated RNA concentration (χ) within hollow structures.

In our system, dsDNA (*χ*) binds to the ERG DBD of FUS-ERG (Fig. 3e(i)). Based on the Nile-red experiments (Supplementary Fig. 5b), which indicate that the ERG DBD exhibits higher hydrophobicity than other regions of FUS-ERG, we hypothesized that dsDNA (*χ*) preferentially diffuses within the hydrophobic regions (*ζ*_*hydrophobic*_) of the hollow co-condensates (**Supplementary Methods**). To test this, we conducted simulations, with results shown in Fig. 4a(iii) and Supplementary Movie 3. The order parameter *ϕ* revealed the gradual formation of an internal surface within the condensates, while the order parameters *χ* and *η* demonstrated the establishment of a hollow condensate structure at equilibrium. Additionally, the distribution of *χ* showed that the simulated dsDNA concentration within the shell region of the condensates (Fig. 4b) closely aligned with the STED experimental data (Fig. 3f(ii)). By varying initial values of *ζ*_*DNA*_ and *ζ*_*protein*_ values, we constructed a phase diagram (Fig. 4c) consistent with the experimental observations (Fig. 1c).

To validate our model, we examined two key parameters (**Supplementary Methods**). The first parameter, ψ_*C*_, was set to 0.85 in the primary simulation (Fig. 4a). When ψ_*C*_ was increased from 0.85 to 0.95, the formation of hollow condensates was not observed (Supplementary Fig. 7a and Supplementary Movie 4). ψ_*C*_ represents the local critical concentration of FUS-ERG required to form the internal surface of hollow condensates with dsDNA containing GGAA microsatellites. A higher ψ_*C*_ indicates that a greater protein concentration is needed to establish the internal surface, reflecting a reduced protein–dsDNA binding affinity. Consistent with this, substituting dsDNA containing GGAA microsatellites with random dsDNA—which exhibits lower binding affinity—also resulted in the absence of hollow co-condensate formation, corroborating the simulation results.

The second parameter, *b*_1_, was set to 0.14 in the main simulation (Fig. 4a). When *b*_1_ was decreased from 0.14 to 0.04, the simulations revealed a failure to form hollow condensates (Supplementary Fig. 7b and Supplementary Movie 5). *b*_1_ quantifies the differential interaction between the hydrophilic component of FUS-ERG and the working buffer relative to its hydrophobic counterpart. A lower *b*_1_ indicates enhanced interaction between the hydrophobic component of FUS-ERG and the working buffer compared to its hydrophilic region. Notably, GFP-FUS-ERG (9RA) (Fig. 1e(iv)) exhibits greater hydrophobicity than wild-type FUS-ERG (Supplementary Fig. 5a). Consistent with this, replacing the wild-type protein with GFP-FUS-ERG (9RA) resulted in the absence of hollow co-condensate formation, corroborating the simulation results.

We next investigated whether our model could replicate the formation of hollow condensates in the PRM-RNA system ^13^ (Fig. 3g) and sought to elucidate the molecular mechanisms distinguishing this system from FUS-ERG hollow condensates. In the PRM-RNA system, RNA interacts with PRM to form a tadpole-like structure. In this configuration, the RNA is oversaturated; one portion binds to PRM, generating hydrophobic regions, while the remaining RNA, due to its hydrophilic nature, contributes to hydrophilic regions within the system. We therefore hypothesized that RNA (*χ*) preferentially diffuses within both the hydrophobic (*ζ*_*hydrophobic*_) and hydrophilic (*ζ*_*hydrophilic*_) regions of the hollow co-condensates (**Supplementary Methods**). To test this hypothesis, we performed simulations (Fig. 4d(i)), with the results shown in Fig. 4d(ii) and Supplementary Movie 6. Unlike the hollow co-condensates of FUS-ERG and dsDNA (Fig. 4b), where dsDNA predominantly localizes on the internal surface, RNA in the PRM-RNA system was distributed across both the inner and outer surfaces of the condensates (Fig. 4e). This distribution closely matched the STED experimental data (Fig. 3g(ii)), revealing a distinct molecular mechanism underlying the formation of hollow condensates in the PRM-RNA system.

Building on the ability of our mathematical model to accurately reproduce the formation of hollow co-condensates, we examined the outcome when FUS-ERG is combined with two types of dsDNA substrates differing in binding affinity. To explore this, we assigned distinct binding affinity coefficients to each dsDNA species, categorizing one as high-affinity (red) dsDNA and the other as low-affinity (blue) dsDNA, while assuming no interactions between the two dsDNA types. Simulations (**Supplementary Methods**) revealed that the low-affinity dsDNA was excluded from the hollow co-condensates (Supplementary Fig. 8a). To experimentally validate this prediction, we first mixed FUS-ERG with dsDNA containing a GGAA tag to form hollow co-condensates (Supplementary Fig. 8b). When non-specific dsDNA was subsequently introduced, it was excluded from the hollow co-condensates, as observed in Supplementary Fig. 8c. These findings highlight the selective nature of hollow co-condensates, driven primarily by protein–dsDNA binding affinity. Interestingly, this inherent property suggests promising potential applications in DNA storage systems, particularly for dynamic data manipulation.

### FET fusion protein-DNA hollow co-condensates enabled dynamic data manipulation for DNA-based information system

In DNA storage, data is encoded in a library of short dsDNA fragments each bearing a unique barcode sequence indexing the encoded content for selective random access. To examine the potential selectivity of hollow co-condensate for specific dsDNA barcodes, we first sought to determine whether hollow co-condensate formation represents a generalizable principle for FET fusion proteins with distinct DBDs and their corresponding dsDNA sequences.

To address this, we introduced a second FET fusion protein, FUS-Gal4, a model system originally developed by McKnight and colleagues ^27^, which we modified by incorporating the FUS RGG motif (amino acids 213–266). After purifying SNAP-FUS- Gal4 (Supplementary Fig. 9a), we observed its ability to form hollow co-condensates with dsDNA containing a single UAS sequence, the specific binding motif for the Gal4 DBD (Fig. 5a(i)). As a control, SNAP-FUS-Gal4 co-localized with dsDNA lacking the UAS sequence in solid condensates (Fig. 5a(ii)). Conversely, we confirmed that SNAP-FUS-ERG forms hollow co-condensates in the presence of GGAA microsatellites but generates solid condensates when GGAA microsatellites are absent (Fig. 5a(iii)-(iv)). These findings demonstrate that the ability of FET fusion proteins to form hollow or solid condensates is determined by the presence of their specific dsDNA sequences.

**Figure 5.**
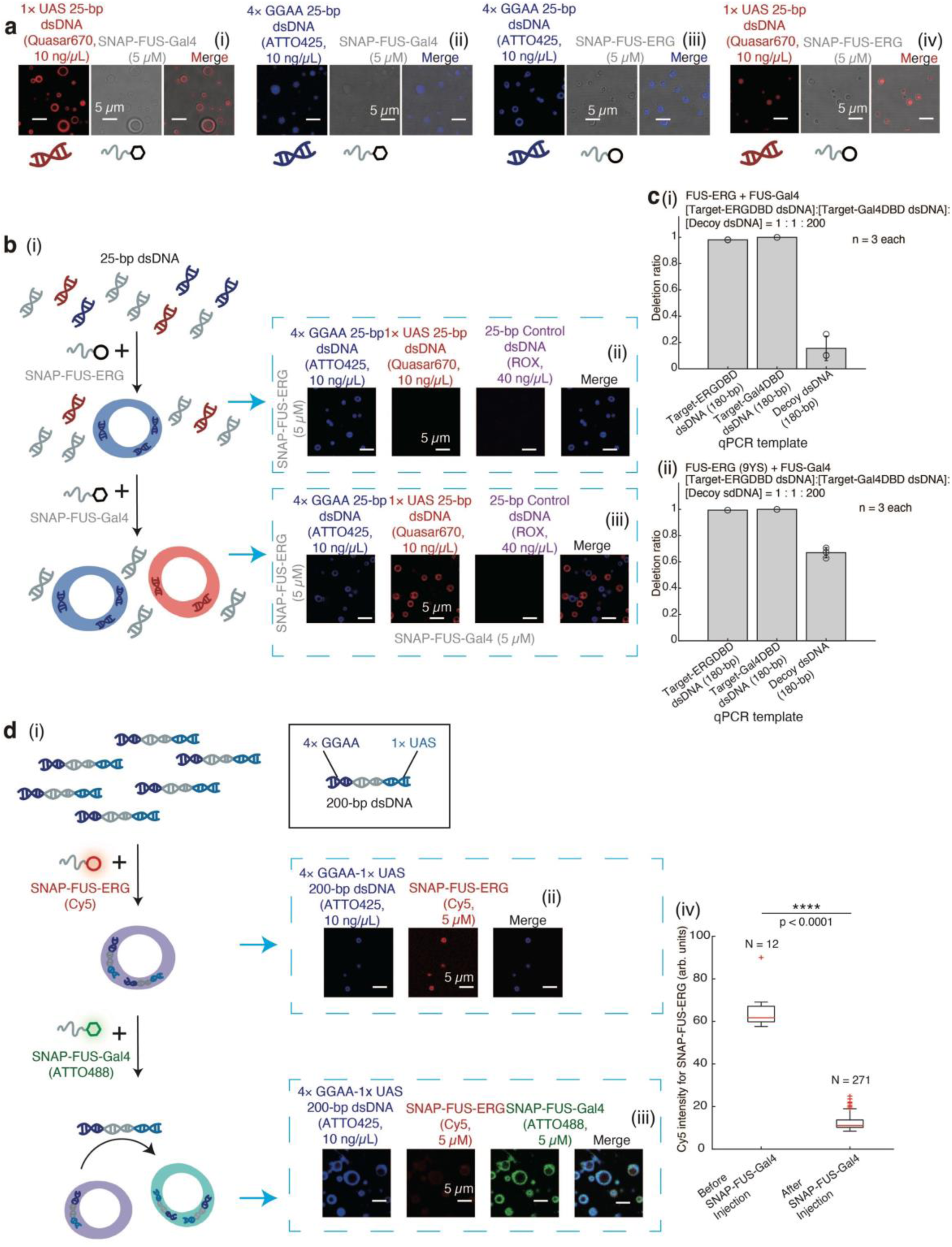
FET fusion protein-DNA hollow co-condensates enabled dynamic data manipulation for DNA-based information system. (**a**) (i)-(ii) 5 μM SNAP-FUS-Gal4 mixed with 10 ng/μL Quasar670-labeled 25-bp 1× UAS dsDNA (target-Gal4DBD dsDNA) (i) or 10 ng/μL (0.6 μM) ATTO425-labeled 25-bp 4× GGAA dsDNA (target-ERGDBD dsDNA) (ii); (iii)-(iv) 5 μM SNAP-FUS-ERG mixed with 10 ng/μL (0.6 μM) ATTO425-labeled target-ERGDBD dsDNA (iii) or 10 ng/μL Quasar670-labeled target-Gal4DBD dsDNA (iv). (**b**) LLPS-based DNA selection. Schematic of the experimental procedure (i). First, 5 μM SNAP-FUS-ERG was added into a dsDNA library containing 10 ng/μL (0.6 μM) ATTO425-labeled target-ERGDBD dsDNA, 10 ng/μL Quasar670- labeled target-Gal4DBD dsDNA, and 40 ng/μL ROX-labeled 25-bp random dsDNA (decoy dsDNA). Fluorescence images were shown in (ii). Second, 5 μM SNAP-FUS- Gal4 was added into the system, and the fluorescence images were shown in (iii). (**c**) Efficiency evaluation of LLPS-based DNA selection (**Methods**). We prepared a dsDNA library containing target-ERGDBD dsDNA, target-Gal4DBD dsDNA, and decoy dsDNA at a molar ratio of 1:1:200. The deletion ratios were measured for these three DNA substrates by qPCR (Supplementary Fig. 11) after the addition of SNAP- FUS-ERG and SNAP-FUS-Gal4 (i) or SNAP-FUS-ERG (9YS) and SNAP-FUS-Gal4 (iii). Independent LLPS-based DNA deletion experiments were repeated three times (n = 3). Error bars, mean ± s.d. (**d**) LLPS-based dynamic and hierarchical data selection. Schematic of the experimental procedure (i). A 180-bp dsDNA substrate containing both 4× GGAA and 1× UAS sequence terminally was synthesized, with the GGAA-end labeled with ATTO425. First, 5 μM Cy5-labeled SNAP-FUS-ERG was mixed with 10 ng/μL these dsDNA substrates. Fluorescence images were shown in (ii). Second, 5 μM Cy5-labeled SNAP-FUS-Gal4 was added, and the fluorescence images were shown in (iii). (iv) Boxplot of Cy5 intensity of SNAP-FUS-ERG in condensates of d(ii)-(iv). The total number N examined over one-time *in vitro* droplet experiments. For the boxplot, the red bar represents median. The bottom edge of the box represents 25^th^ percentiles, and the top is 75^th^ percentiles. Most extreme data points are covered by the whiskers except outliers. The ‘+’ symbol is used to represent the outliers. Statistical significance was analyzed using unpaired t test for two groups. P value: two-tailed; p value style: GP: 0.1234 (ns), 0.0332 (*), 0.0021 (**), 0.0002 (***), <0.0001 (****). Confidence level: 95%. The working buffer containing 40 mM Tris-HCl (pH 7.5), 150 mM KCl, 2 mM MgCl_2_, 1 mM DTT and 0.2 mg/mL BSA.

Leveraging the two orthogonal DBD-dsDNA interactions, we first explored whether hollow co-condensates could be utilized to sort specific dsDNA substrates from a dsDNA library. We mixed three dsDNA substrates in a 1:1:4 ratio of 25-bp 4× GGAA dsDNA (target-ERGDBD dsDNA), 25-bp dsDNA containing a 1× UAS sequence (target-Gal4DBD dsDNA), and 25-bp random dsDNA (decoy dsDNA, Fig. 5b(i)). Initially, we added SNAP-FUS-ERG into the dsDNA library to observe blue-colored hollow co-condensates (Fig. 5b(ii)), affirming that SNAP-FUS-ERG selectively binds to target-ERGDBD dsDNA, while target-Gal4DBD dsDNA and decoy dsDNA remain in the solvent phase. Building upon this observation, when we subsequently introduced SNAP-FUS-Gal4, besides the blue-colored hollow condensates, a secondary set of hollow co-condensates emerged exhibiting a red hue (Fig. 5b(iii)), indicating the spatial sorting of dsDNA into separate condensates according to their distinct barcodes.

We further demonstrate through control experiments that the hollow co-condensate architecture is essential for selective dsDNA sorting. When GFP-FUS-ERG (9YS) was mixed with target-ERGDBD dsDNA or target-Gal4DBD dsDNA (used as control dsDNA), no hollow co-condensates formed (Supplementary Fig. 10a and Fig. 1e(iv)). Repeating the experiment shown in Fig. 5b, but substituting the initial addition with GFP-FUS-ERG (9YS), revealed that this variant not only failed to selectively bind to target-ERGDBD dsDNA (Supplementary Fig. 10b(i)), but also disrupted the subsequent sorting of target-Gal4DBD dsDNA by SNAP-FUS-Gal4 (Supplementary Fig. 10b(ii)). Additionally, the process significantly affected decoy dsDNA substrates, ultimately resulting in a complete failure of DNA deletion.

These observations strongly imply that hollow co-condensates could be harnessed for targeted DNA deletion within DNA libraries. Crucially, we also examined whether the non-specific dsDNA substrates in the library remained unaffected during this process. To evaluate selective DNA deletion efficiency, we prepared a dsDNA library containing target-ERGDBD dsDNA, target-Gal4DBD dsDNA, and decoy dsDNA at a molar ratio of 1:1:200. We independently used GFP-FUS-ERG and SNAP-FUS-Gal4 to sequester and pull-down target-ERGDBD and target-Gal4DBD dsDNA, respectively, by iterative centrifugation (**Methods**). After four iterations, quantitative real-time PCR (qPCR) revealed that target-ERGDBD dsDNA and target-Gal4DBD dsDNA was deleted with an efficiency of 98.1 ± 0.1% and 100 ± 0.0%, respectively, while the decoy dsDNA was removed at a much lower rate of 15.6 ± 9.3% (Fig. 5c(i)) according to the calibration of each dsDNA in the library (Supplementary Fig. 11). As a control, we repeated the experiment using the mutant GFP-FUS-ERG (9YS) instead of the wild-type GFP-FUS-ERG. Although target-ERGDBD dsDNA and target-Gal4DBD dsDNA were deleted at similar efficiencies of 99.4 ± 0.0% and 100 ± 0.0%, respectively, decoy dsDNA was strongly depleted by 67.0 ± 3.9%, a 4.3-fold excess removal compared to deletion by the wildtype FUS-ERG (Fig. 5c(ii)).

Lastly, we challenged the hollow co-condensate system for dynamic and hierarchical data selection. More specifically, we implemented a two-step sequential sorting logic based on the barcode-protein binding specificity (Fig. 5d(i)). A 180-bp dsDNA substrate containing both 4× GGAA and 1× UAS sequence terminally was synthesized, with the GGAA-end labeled with ATTO425. Initially, we mixed the dsDNA substrate with SNAP-FUS-ERG and observed hollow co-condensates (Fig. 5d(ii)), indicating the binding of SNAP-FUS-ERG to the 4× GGAA sequence. Afterwards, SNAP-FUS-Gal4 was introduced into the sample. We found that SNAP-FUS-Gal4 displaced SNAP-FUS-ERG from the condensates, forming new hollow co-condensates with the dsDNA substrates (Fig. 5d(iii)). This outcome suggests that SNAP-FUS-Gal4 effectively competes with SNAP-FUS-ERG for binding to the dsDNA substrates and as a result, the FUS-ERG condensate disassembled (Fig. 5d(iv)). In stark contrast, with the order of addition switched to SNAP-FUS-Gal4 in the first step followed by SNAP-FUS-ERG in the second, all components co-localized into homogeneous condensates (Supplementary Fig. 12a). Considering the stronger binding affinity of Gal4 DBD to UAS sequence compared to ERG DBD to GGAA microsatellites (Supplementary Fig. 9a(iii) and 1b(iii)), the dynamic sequential logic of data manipulation necessitates the strict temporal programming of hollow co-condensate system.

As control experiments, we evaluated several variant FET fusion proteins, including FUS-Gal4 lacking the RGG motif (GFP-FUS-Gal4 (no RGG), Supplementary Fig. 9b), GFP-FUS-ERG (9YS) (Supplementary Fig. 1c), and GFP-FUS-ERG (9RA) (Supplementary Fig. 1f). Independent assessments confirmed that these variants were unable to drive hollow co-condensate formation, rather, they formed homogeneous condensates with target dsDNA (Supplementary Fig. 12b, d(i), and e(i) & Fig. 1d(ii) and e(iv)). The experiment depicted in Fig. 5d was repeated, substituting the wildtype fusion proteins with their mutants (Supplementary Fig. 12c-e). Despite maintaining the correct order of protein addition, target selection failed at the step where the mutant proteins were introduced. These findings underscore the critical requirement for both FET fusion proteins to possess the intrinsic capability to form hollow co-condensates in order to achieve the outcomes observed in Fig. 5d.

## Discussion

In this study, we demonstrate that the FET family fusion oncoprotein FUS-ERG forms hollow co-condensates with dsDNA containing GGAA microsatellites. Using biochemical assays, super-resolution imaging, and mathematical modeling, we elucidate the molecular mechanism as nested asymmetric phase separation. Moreover, we show that the self-organization and morphology control of FUS-ERG hollow co-condensates can be leveraged for DNA-based information manipulation, enabling precise DNA deletion within dsDNA libraries and supporting dynamic, hierarchical data selection.

Biomolecular condensates can be modeled using three primary approaches: microscopic, mesoscopic, and macroscopic models. Microscopic models, such as molecular dynamics and coarse-grained simulations, provide detailed insights into the structural and dynamic properties of condensate formation. This approach has been successfully applied to hollow co-condensates formed by PRM and RNA ^13^. However, the high computational demands of microscopic models make them impractical for our system. Macroscopic models, including phase-field models based on Flory-Huggins theory ^12,28^, effectively capture component interactions and macroscopic phase structures within condensates. Despite these strengths, macroscopic models fall short in resolving local characteristics within the shell regions of hollow co-condensates, such as hydrophobic distributions and fine-scale component organization—features that are critical for our study. To overcome these limitations, we adopted a molecularly informed phase-field model ^24^, which operates at the mesoscopic scale. This model incorporates order parameters *ϕ* and *χ* (Fig. 4), enabling a seamless integration of macroscopic phase structures with local molecular characteristics. Compared to microscopic and macroscopic approaches, this mesoscopic model offers a balanced and computationally feasible strategy to explore the complex behavior of hollow co-condensates in our system (**Supplementary Methods**).

Despite our observation of dsDNA containing GGAA microsatellites translocating into condensates (Fig. 2), the molecular mechanism underlying this translocation remains elusive. To address this, we propose an interesting hypothesis: Initially, Closed state proteins undergo LLPS with the working buffer, forming a condensed phase of Closed state proteins within solid condensates. Upon binding of dsDNA substrates containing GGAA motifs, conformational changes are induced, resulting in the formation of Open-state protein–dsDNA complexes. Due to their relative scarcity, these Open-state complexes establish a loosened phase ^29^. These Open-state protein–dsDNA complexes segregate from the condensed phase of Closed state proteins, creating microscopic channels within the condensates. As additional dsDNA substrates bind to FUS-ERG molecules on the condensate surface, these channels facilitate the diffusion of Open-state complexes into the condensate interior. This process is likely driven by a combination of interfacial depletion effects ^30,31^ and hydrophobic interactions. As the Open-state complexes diffuse inward, they displace Closed state proteins, establishing an inner interface enriched with Open-state complexes. Collectively, this hypothesis suggests that hydrophobic interactions and interfacial depletion effects synergistically drive the internal phase separation of Open-state and Close-state proteins, ultimately leading to the formation of diffusion-controlled, hollow co-condensates. Further investigation into this proposed mechanism represents an exciting avenue for future research.

Deciphering the molecular mechanisms of condensate architectures has important biological relevance in understanding the living cells, which is one of important properties of multicomponent phase-separated cellular bodies. For instance, in mammalian nucleoli, dense fibrillar components (DFCs) are one type of biomolecular condensates, which envelop another type of biomolecular condensates – fibrillar centers (FCs), creating “Russian doll” structures that facilitate efficient transcription at the FC/DFC interface ^32^. The second example is during *in vitro* reconstitution of mitochondrial transcription machinery, the formation of hollow co-condensates in mitochondrial machinery plays a crucial role in regulating transcription rates ^14^. Thirdly, FET family fusion oncoproteins have been found to form biomolecular condensates at genomic binding sites ^33,34^, and these condensates can recruit RNA polymerase II to regulate gene transcription, causing sarcomas and leukemia ^35,36^. Whether the condensate architectures can regulate gene transcription is one interesting topic for the future study.

Finally, understanding the molecular mechanism of hollow co-condensate also provide potential applications in biotechnology, especially in DNA-based information manipulation. We reconstitute hollow condensates, a natural mode of spatial regulation, in synthetic DNA information systems, and showed that this specific architecture is necessary for highly selective and highly dynamic data manipulation. Although DNA self-assembly has been successfully applied to the field of DNA-based data storage ^37–39^, we provide the first direct demonstration that protein-DNA self-assembly and biomolecular condensate morphology control can be used for spatial information manipulation.

For the targeted DNA deletion within DNA libraries (Fig. 5b), we compared the separation efficiency of our method with previous methods. So far, data sorting has been demonstrated by using complementary DNA probes combined with solid state purification or site-specific restriction enzyme cleavage of a particular DNA from an adsorption matrix ^37,40^. In particular, primer-based magnetic separation was shown to have < 75% efficiency with a 20-nt primer length and optimal temperature ^37^. Restriction enzyme cleavage was shown to have around 40% of target-strand retention ^41^. Our LLPS-based manipulation, in comparison, achieved almost 100% removal while affecting only 15% of decoy DNA in an oligo system of notably lower signal-to-noise ratio (1:1:200, Fig. 5c). Spatial regulation by protein-DNA assembly, compared to widely used DNA self-assembly approaches, could leverage several unique properties for the observed high sorting efficiency and accuracy. First, complex protein-DNA interactions may lead to enhanced binding efficiencies and selectivity. Moreover, they could offer more broad and tunable affinity profiles. Second, the condensate morphology induced by protein conformational switch and the heterogeneity of internal DNA distribution may have further enhanced the condensate’s target selectivity (Supplementary Fig. 8a).

On the other hand, the results on hierarchical data sorting (Fig. 5d) suggested that the hollow co-condensates are also sufficiently dynamic for compositional exchange. Therefore, the architecture acts as a temporary storage depot for DNA molecules, allowing for sequential manipulations of data encoded in DNA, while ensuring a high indexing specificity to the programmed temporal logics. Therefore, our LLPS-based hollow co-condensate system represents a new vessel for in-storage dynamic regulation and data processing in memory. Building upon this basic principle, it is possible to include more dsDNA-DBD binding for finer information sorting, to employ more complex architectures for precise and flexible multi-component control, and to enable sophisticated information manipulation through protein engineering and protein-based regulation.

In summary, these findings provide important insights into the biophysical mechanisms underlying multicomponent phase-separated cellular bodies, and also offer innovative strategies for manipulating DNA-based information.

## Author Contributions

L.Z. prepared biological samples, designed and conducted all experiments, performed data analysis, and wrote the manuscript. Q.G. and L.Z. built up the mathematical model, performed simulation and theoretical analysis, and wrote the manuscript. K.Z. and Z.C. conducted the super-resolution imaging experiments. B.J. assisted L.Z. for the qPCR experiments. Y.X. supervised the project, experimental designs, and wrote the manuscript. L.Q. designed the experiments for DNA manipulation, and wrote the manuscript. Z.Q. supervised the project, experimental designs, and data analysis, and wrote the manuscript with input from all authors.

## Acknowledgments

We thank the Peking Nanofab for process support. We thank the contributions of the Engineering Research Center for Semiconductor Integrated Technology, Institute of Semiconductors, Chinese Academy of Sciences. We thank Dr. Jun Cheng for assisting us for protein purification and *in vivo* experiments. We thank Yijia Bian (Undergraduate in the Integrated Science Program, Peking University) for cartoon figures plotting. We thank Dr. Luhua Lai, Dr. Chun Tang, Dr. Yujie Sun, and Dr. Zhiyuan Li (Peking University), Dr. Wei Ji (Institute of Biophysics, Chinese Academy of Sciences), and the members of the Z.Q. laboratory for comments on the manuscript.

## References

1 Li, C., Guo, M.-T., He, X., Liu, Q.-X. & Qi, Z. Biomolecular condensates bridge experiment and theory of mass-conserving reaction-diffusion systems in phase separation. bioRxiv, 2024.2008.2008.607271, doi:10.1101/2024.08.08.607271 (2024).

2 Banani, S. F., Lee, H. O., Hyman, A. A. & Rosen, M. K. Biomolecular condensates: organizers of cellular biochemistry. Nat Rev Mol Cell Bio 18, 285–298, doi:10.1038/nrm.2017.7 (2017).

3 Lyon, A. S., Peeples, W. B. & Rosen, M. K. A framework for understanding the functions of biomolecular condensates across scales. Nat Rev Mol Cell Bio 22, 215–235, doi:10.1038/s41580-020-00303-z (2021).

4 Cardona, A. H. et al. Self-demixing of mRNA copies buffers mRNA:mRNA and mRNA:regulator stoichiometries. Cell 186, 4310-+, doi:10.1016/j.cell.2023.08.018 (2023).

5 Alberti, S. & Dormann, D. Liquid-Liquid Phase Separation in Disease. Annual Review of Genetics*, Vol* 53 **53**, 171–194, doi:10.1146/annurev-genet-112618-043527 (2019).

6 Li, C. et al. Deciphering the molecular mechanism underlying morphology transition in two-component DNA-protein cophase separation. Structure 33, 62–77.e68, doi:10.1016/j.str.2024.10.026 (2025).

7 Alshareedah, I., Moosa, M. M., Pham, M., Potoyan, D. A. & Banerjee, P. R. Programmable viscoelasticity in protein-RNA condensates with disordered sticker-spacer polypeptides. Nature Communications 12, 6620, doi:10.1038/s41467-021-26733-7 (2021).

8 Boeynaems, S. et al. Spontaneous driving forces give rise to protein-RNA condensates with coexisting phases and complex material properties. P Natl Acad Sci USA 116, 7889–7898, doi:10.1073/pnas.1821038116 (2019).

9 Ma, W., Zhen, G., Xie, W. & Mayr, C. In vivo reconstitution finds multivalent RNA–RNA interactions as drivers of mesh-like condensates. eLife 10, e64252, doi:10.7554/eLife.64252 (2021).

10 Mao, S., Kuldinow, D., Haataja, M. P. & Kosmrlj, A. Phase behavior and morphology of multicomponent liquid mixtures. Soft Matter 15, 1297–1311, doi:10.1039/c8sm02045k (2019).

11 Shin, Y. & Brangwynne, C. P. Liquid phase condensation in cell physiology and disease. Science 357, doi:ARTN eaaf4382 10.1126/science.aaf4382 (2017).

12 Yu, H. et al. HSP70 chaperones RNA-free TDP-43 into anisotropic intranuclear liquid spherical shells. 371, eabb4309, doi:10.1126/science.abb4309 %J Science (2021).

13 Alshareedah, I., Moosa, M. M., Raju, M., Potoyan, D. A. & Banerjee, P. R. Phase transition of RNA-protein complexes into ordered hollow condensates. P Natl Acad Sci USA 117, 15650–15658, doi:10.1073/pnas.1922365117 (2020).

14 Feric, M. et al. Mesoscale structure–function relationships in mitochondrial transcriptional condensates. Proceedings of the National Academy of Sciences 119, e2207303119, doi:doi:10.1073/pnas.2207303119 (2022).

15 Sizemore, G. M., Pitarresi, J. R., Balakrishnan, S. & Ostrowski, M. C. The ETS family of oncogenic transcription factors in solid tumours. Nature Reviews Cancer 17, 337–351, doi:10.1038/nrc.2017.20 (2017).

16 Shing, D. C. et al. FUS/ERG gene fusions in Ewing’s tumors. Cancer Research 63, 4568–4576 (2003).

17 Guo, H. T. et al. Spatial engineering of with addressable phase-separated RNAs. Cell 185, 3823-+, doi:10.1016/j.cell.2022.09.016 (2022).

18 Fei, Z. J., Gupta, N., Li, M. J., Xiao, P. F. & Hu, X. Toward highly effective loading of DNA in hydrogels for high-density and long-term information storage. Sci Adv 9, doi:ARTN eadg9933, 10.1126/sciadv.adg9933 (2023).

19 Boegels, B. W. A. et al. DNA storage in thermoresponsive microcapsules for repeated random multiplexed data access. Nat Nanotechnol 18, 912-+, doi:10.1038/s41565-023-01377-4 (2023).

20 Kato, M. et al. Cell-free Formation of RNA Granules: Low Complexity Sequence Domains Form Dynamic Fibers within Hydrogels. Cell 149, 753–767, doi:10.1016/j.cell.2012.04.017 (2012).

21 Wang, J. et al. A Molecular Grammar Governing the Driving Forces for Phase Separation of Prion-like RNA Binding Proteins. Cell 174, 688–699, doi:10.1016/j.cell.2018.06.006 (2018).

22 Davis, R. B., Kaur, T., Moosa, M. M. & Banerjee, P. R. FUS oncofusion protein condensates recruit mSWI/SNF chromatin remodeler via heterotypic interactions between prion-like domains. Protein Sci 30, 1454–1466, doi:10.1002/pro.4127 (2021).

23 Lukinavičius, G. et al. Stimulated emission depletion microscopy. Nature Reviews Methods Primers 4, 56, doi:10.1038/s43586-024-00335-1 (2024).

24 Han, Y. C., Xu, Z. R., Shi, A. C. & Zhang, L. Pathways connecting two opposed bilayers with a fusion pore: a molecularly-informed phase field approach. Soft Matter 16, 366–374, doi:10.1039/c9sm01983a (2020).

25 Ito, A. Domain patterns in copolymer-homopolymer mixtures. Phys Rev E 58, 6158–6165, doi:DOI 10.1103/PhysRevE.58.6158 (1998).

26 Ohta, T. & Ito, A. Dynamics of Phase-Separation in Copolymer-Homopolymer Mixtures. Phys Rev E 52, 5250–5260, doi:DOI 10.1103/PhysRevE.52.5250 (1995).

27 Kwon, I. et al. Phosphorylation-Regulated Binding of RNA Polymerase II to Fibrous Polymers of Low-Complexity Domains. Cell 155, 1049–1060, doi:10.1016/j.cell.2013.10.033 (2013).

28 Shin, Y. et al. Spatiotemporal Control of Intracellular Phase Transitions Using Light-Activated optoDroplets. Cell 168, 159-+, doi:10.1016/j.cell.2016.11.054 (2017).

29 Regan, M. C. et al. Structural and dynamic studies of the transcription factor ERG reveal DNA binding is allosterically autoinhibited. P Natl Acad Sci USA 110, 13374–13379, doi:10.1073/pnas.1301726110 (2013).

30 Kim, K. et al. Processable high internal phase Pickering emulsions using depletion attraction. Nature Communications 8, 14305, doi:10.1038/ncomms14305 (2017).

31 Ji, S. & Walz, J. Y. Depletion forces and flocculation with surfactants, polymers and particles — Synergistic effects. Current Opinion in Colloid & Interface Science 20, 39–45, 10.1016/j.cocis.2014.11.006 (2015).

32 Yao, R.-W. et al. Nascent Pre-rRNA Sorting via Phase Separation Drives the Assembly of Dense Fibrillar Components in the Human Nucleolus. Mol Cell 76, 767–783.e711, doi:10.1016/j.molcel.2019.08.014 (2019).

33 Chong, S. S. et al. Imaging dynamic and selective low-complexity domain interactions that control gene transcription. Science 361, 378-+, doi:10.1126/science.aar2555;eaar2555 (2018).

34 Zuo, L. et al. Loci-specific phase separation of FET fusion oncoproteins promotes gene transcription. Nature Communications 12, 1491, doi:10.1038/s41467-021-21690-7 (2021).

35 Thomsen, C., Grundevik, P., Elias, P., Stahlberg, A. & Aman, P. A conserved N-terminal motif is required for complex formation between FUS, EWSR1, TAF15 and their oncogenic fusion proteins. Faseb Journal 27, 4965–4974, doi:10.1096/fj.13-234435 (2013).

36 Ichikawa, H., Shimizu, K., Hayashi, Y. & Ohki, M. An Rna-Binding Protein Gene, Tls/Fus, Is Fused to Erg in Human Myeloid-Leukemia with T(16,21) Chromosomal Translocation. Cancer Research 54, 2865–2868 (1994).

37 Lin, K. N., Volkel, K., Tuck, J. M. & Keung, A. J. Dynamic and scalable DNA-based information storage. Nature Communications 11, doi:10.1038/s41467-020-16797-2 (2020).

38 Chen, K. K., Zhu, J. B., Boskovic, F. & Keyser, U. F. Nanopore-Based DNA Hard Drives for Rewritable and Secure Data Storage. Nano Lett 20, 3754–3760, doi:10.1021/acs.nanolett.0c00755 (2020).

39 Zhang, Y. N. et al. DNA origami cryptography for secure communication. Nature Communications 10, doi:ARTN 5469 10.1038/s41467-019-13517-3 (2019).

40 Tomek, K. J. et al. Driving the Scalability of DNA-Based Information Storage Systems. Acs Synth Biol 8, 1241–1248, doi:10.1021/acssynbio.9b00100 (2019).

41 Lin, K. N. et al. A primordial DNA store and compute engine. Nat Nanotechnol, doi:10.1038/s41565-024-01771-6 (2024).

